# A Dual Inhibitory Network in the Thalamic Reticular Nucleus Delineated by Pallidal and Intra-Reticular Inhibition

**DOI:** 10.1101/2025.10.05.680571

**Authors:** Frances S. Cho, John R. Huguenard, Jeanne T. Paz

**Affiliations:** Gladstone Institute of Neurological Disease, Gladstone Institutes, San Francisco CA 94158; Department of Neurology, University of California San Francisco, San Francisco CA 94158; Kavli Institute for Fundamental Neuroscience, University of California San Francisco, San Francisco CA 94158; Department of Neurology & Neurological Sciences, Stanford University School of Medicine, Stanford CA 94305

**Keywords:** Thalamic Reticular Nucleus (TRN), External Globus Pallidus (GPe), GABAergic inhibition, Inhibitory Circuit Motifs, Counter-Inhibition and Feed-Forward Circuit Motifs

## Abstract

Long described as an inhibitory “guardian of the gateway,” the thalamic reticular nucleus (TRN) shapes which thalamic signals reach the cortex during attention, arousal, and sensory processing. However, how the inhibitory wiring within TRN supports this flexible gating—from modality-specific tuning to global control—remains poorly defined. Using cell type–specific optogenetic input mapping and whole-cell patch-clamp recordings in mice, we dissect inhibitory connectivity in TRN from two major GABAergic sources: the TRN itself and the external globus pallidus (GPe). All recorded TRN neurons received inhibition from the GPe, whereas a subset also received intra-TRN inhibition. Intra-TRN inhibition arose predominantly from somatostatin-expressing onto parvalbumin-expressing TRN neurons (SOM→PV), revealing subtype-specific connectivity recruited by thalamic excitation to form a feedforward motif. These findings delineate a dual inhibitory architecture: local intra-TRN circuits provide spatially selective inhibition, whereas pallidal inputs deliver diffuse inhibition. These complementary mechanisms may support both modality-specific gating and global state-dependent control of thalamic output.

**Graphical Abstract:** The thalamic reticular nucleus (TRN) integrates two distinct inhibitory motifs. Sparse, dendrite-targeting intra-TRN connections from parvalbumin- and somatostatin-expressing neurons provide structured, feedforward inhibition recruited by thalamocortical (TC) inputs. In parallel, robust and widespread inhibition from Npas1-expressing neurons of the external globus pallidus (GPe) supplies a strong external drive that can coordinate activity across the TRN. Together, these complementary motifs define a dual inhibitory architecture that may shape thalamocortical gating and highlight cellular entry points for therapeutic intervention.

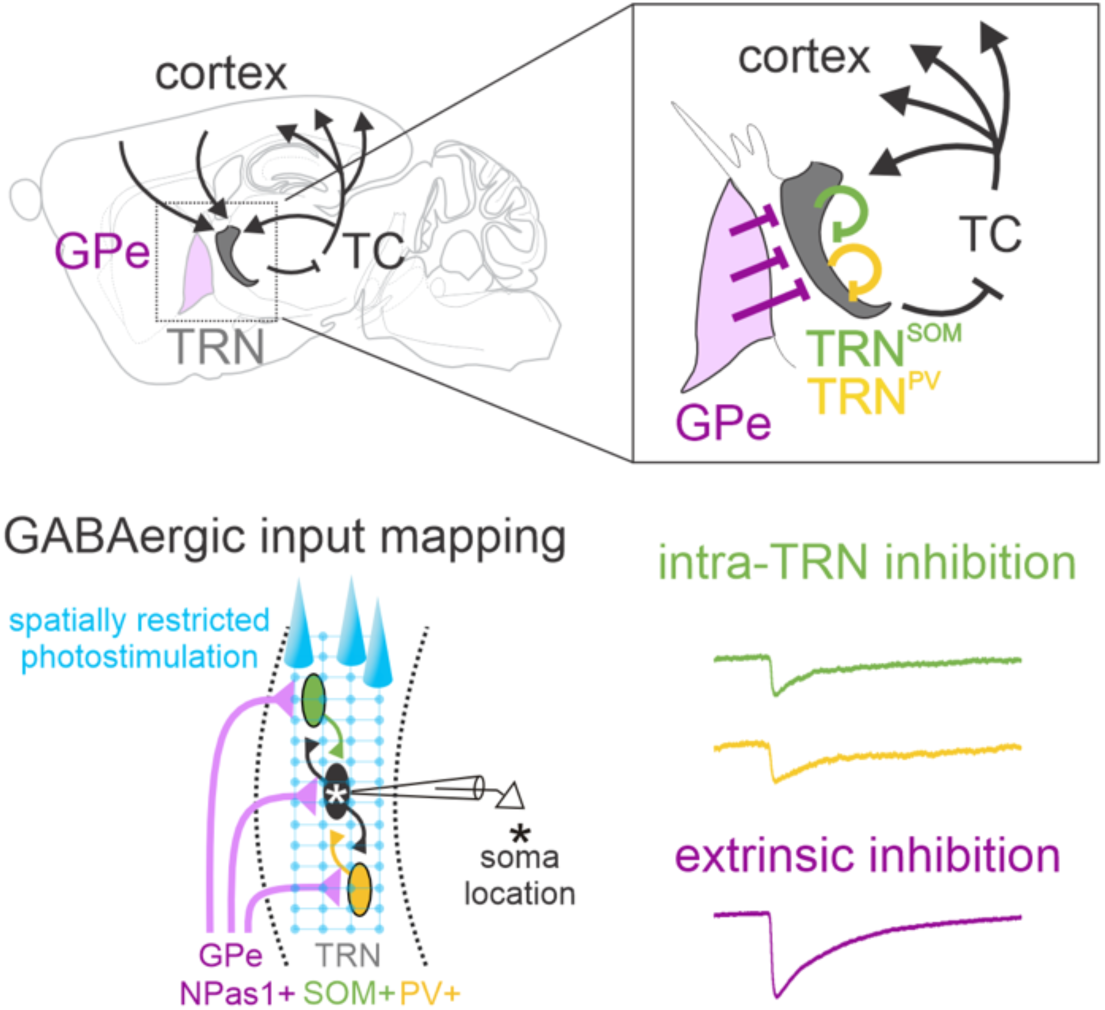

## Introduction

The thalamic reticular nucleus (TRN) is a central inhibitory hub that gates thalamocortical communication and regulates attention, arousal, and sensory processing^1–5^. Comprised entirely of GABAergic neurons, the TRN exerts powerful inhibition onto thalamic relay nuclei, yet the organization of inhibition within the TRN itself remains poorly understood^6^. The TRN contains molecularly and functionally distinct subnetworks with specialized input–output connectivity^3,7–11^. Retrograde rabies tracing identified the external globus pallidus (GPe) and the TRN itself as major GABAergic sources of input^8^.

While GPe projections to TRN have been recognized for decades^12^, their function has remained largely unexplored. Similarly, whether TRN neurons form chemical inhibitory synapses with one another is debated (reviewed in ^4,13^). Gap junction-coupled intra-TRN electrical synapses have been consistently demonstrated by multiple groups and suggested to play a role in synchronizing TRN neurons^14–17^. However, the existence of intra-TRN chemical GABAergic synapses has been a subject of controversy, with multiple electrophysiological studies suggesting their existence and functionality in ferrets, rats, and mice^16,18,19^, and others failing to find them^20–22^. This controversy highlights the need for systematic analysis of TRN inhibitory connectivity across molecularly defined subtypes and comparison with extrinsic pallidal input. Understanding how these inhibitory motifs are organized is critical for uncovering mechanisms of thalamocortical synchrony and its disruption in conditions such as absence epilepsy, schizophrenia, and attention disorders.

Here, we dissect inhibitory connectivity within the TRN and compare it with extrinsic inputs from the GPe. Using whole-cell recordings combined with cell type–specific optogenetic input mapping in adult mice, we show that TRN neurons receive sparse, spatially organized inhibition that is mainly, but not exclusively, driven by somatostatin-expressing TRN neurons, alongside robust extrinsic input from Npas1-expressing GPe neurons. Together, these inputs define a dual inhibitory architecture that complements electrical coupling within TRN, revealing circuit motifs that may regulate thalamocortical synchrony and provide cellular entry points for therapeutic modulation in neuropsychiatric and seizure disorders.

## Results

To compare extrinsic and intrinsic sources of GABAergic inhibition in TRN, we examined molecularly distinct sources of presynaptic input providing GABAergic inhibition to the TRN. We targeted GPe inputs because they represent a major source of GABAergic inhibition to TRN, with Npas1-positive GPe (GPe^Npas^) neurons recently shown to form functional connections in the TRN^23^. To assess intra-TRN inhibition, we examined whether somatostatin- or parvalbumin-expressing TRN neurons (TRN^SOM^ and TRN^PV^), the principal molecularly defined TRN subpopulations^8,9,24–27^, are the main sources of intra-TRN inhibition. Cre-dependent AAV-EF1α-hChR2 was injected into the GPe of Npas1-Cre mice or into the TRN of Parvalbumin-Cre or Somatostatin-Cre mice. Opsin expression was allowed to develop for six weeks to ensure robust expression in axons. All experiments were performed in adult mice (2–7 months old). Monosynaptic inputs to TRN neurons were mapped using focal (2 μm beam) photostimulation with a 460 × 100 μm grid spanning the anterior-posterior axis of the TRN (**Figure 1A, Supp. Figure 1A**). Evoked synaptic responses were easily separable from any direct photocurrent based on latency and were abolished in the presence of the GABA_A_R antagonist gabazine (see Methods, **Supp. Figure 1**). Response maps were constructed by quantifying only the evoked synaptic responses at sites with reliably evoked responses across multiple trials. This approach enabled activation of defined GABAergic populations and mapping of their synaptic inputs onto TRN neurons (**Figure 1A, Supp. Figure 1**).

**Figure 1.**
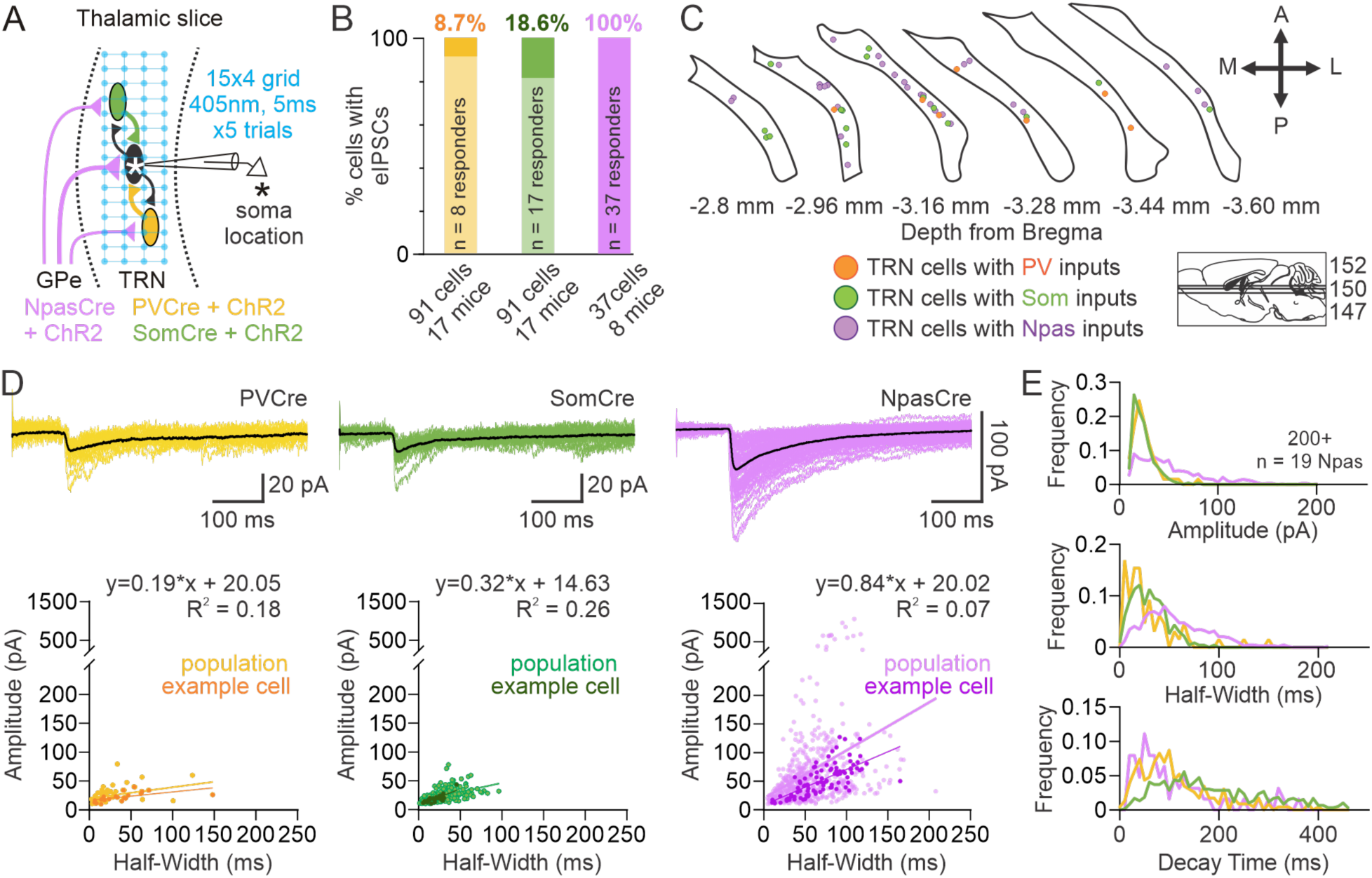
TRN neurons receive GABAergic synaptic inputs from intra-TRN and external globus pallidus. **A.** Schematic of focal photostimulation. AAV2/5-DIO-ChR2-eYFP was injected into the TRN of PV-Cre or Som-Cre mice, and into the GPe of Npas-Cre mice. Whole-cell voltage-clamp recordings were obtained from TRN neurons while ChR2-expressing axons were focally stimulated in a 15 × 4 grid (spanning 460 × 100 µm) along the anterior-posterior and medial-lateral axes of the TRN. The asterisk * indicates the location of the recorded cell body. At each site, a 5 ms, 405 nm laser pulse was delivered. Stimulation sites were activated in randomized order at 2 Hz (one site every 0.5 s). **B.** Proportion of TRN neurons showing evoked inhibitory postsynaptic currents (eIPSCs) in response to activation of TRN^PV^, TRN^SOM^, or GPe^NPas^ inputs. n = 8 of 91 cells from 17 PV-Cre mice; n = 17 of 91 cells from 17 Som*-*Cre mice; n = 37 of 37 cells from 8 Npas*-*Cre mice. **C.** Location of postsynaptic TRN cells responding to TRN^PV^, TRN^SOM^, or GPe^Npas^ inputs in the dorsal-ventral horizontal plane. Brain schematic adapted from ^45^. **D.** Representative TRN eIPSCs recorded at –70 mV in response to optogenetic activation of PV, Som, or Npas synaptic inputs. Black traces indicate averages of all synaptic events from each cell; gray traces show individual events. Scatter plots (bottom) show eIPSC’s half-width (time to 50% decay from peak) versus amplitude. Each point represents an individual eIPSC. PV-triggered eIPSCs: n = 65 events from 4 cells; Som*-*triggered eIPSCs: n = 232 events from 13 cells; Npas-triggered eIPSCs: n = 635 events from 15 cells. **E.** Relative frequency histograms of eIPSC amplitude (5 pA bins), half-width (5 ms bins), and decay time (10 ms bins).

### Prevalence of Synaptic Inhibition within TRN

The prevalence of responses varied sharply depending on the source of inhibition. Optical activation of GPe^Npas^ axons in the TRN elicited inhibitory postsynaptic currents (IPSCs) in all recorded TRN neurons (37/37 cells from 8 mice) (**Figure 1B**). In contrast, optical activation of TRN^SOM^ neurons evoked IPSCs in 18.6% of TRN neurons (17/91 cells from 17 mice), and activation of TRN^PV^ neurons evoked IPSCs in 8.7% (8/91 cells from 17 mice) (**Figure 1B**). Because the axons in the TRN may be sparse and thin, we next investigated whether a spatially larger illumination of the weakest (i.e., lowest-prevalence) presynaptic input could unmask larger intra-TRN connectivity. Therefore, we stimulated TRN^PV^ neurons using wider-field illumination with a 200-μm optical fiber (**Supp. Figure 2A**) as opposed to a focal 2 μm beam (**Figure 1A**). This wider illumination evoked monosynaptic IPSCs in 46.4% of TRN neurons (13 of 28, **Supp. Figure 2B–C**), which we observed consistently with two distinct opsins, ChR2 (5 of 15 cells) and CheTA (8 of 14) (pooled data, **Supp. Figure 2B**). The kinetics of these IPSCs were consistent with prior reports and matched those observed in our focal mapping experiments (**Supp. Figure 2D–E**). In sum, the wider illumination revealed 5 times more connectivity than the focal mapping, suggesting that the intra-TRN connections are sparse and likely originate from distal locations. Altogether, GPe^Npas^ inputs robustly inhibited all TRN neurons tested, while recurrent intra-TRN synaptic inhibition originating from TRN^SOM^ or TRN^PV^ neurons was sparse and likely distal.

### Location of TRN neurons receiving GABAergic inhibition

Because the TRN can be functionally divided into various modalities and sectors, we next asked whether the TRN neurons that receive intra-TRN versus GPe→TRN inputs are located in any specific region. TRN neurons receiving inhibition from GPe^Npas^, TRN^PV^, or TRN^SOM^ cellular sources were observed at every level of what was sampled along the dorsal-ventral axis. Furthermore, TRN neurons receiving inhibition were found across the entire anterior-posterior axis of the TRN (**Figure 1C**). We next characterized the distribution of TRN neurons receiving inhibition across the medio–lateral axis of the TRN by classifying the location into the TRN “core” or “shell” (with “core” defined as the central 50% along the medial–lateral axis and “shell” defined as the medial and lateral 25%, similar to previous studies^10^). Interestingly, we found that intra-TRN inputs (from TRN^PV^ or TRN^SOM^) were enriched in the TRN “core” compared to the TRN “shell,” with 16.8% of all cells recorded in the “core” (23 of 137 cells) exhibiting eIPSCs in response to photostimulation of TRN^PV^ or TRN^SOM^ inputs, compared to 3.2% prevalence of intra-TRN recipients in the “shell” (1 of 31 cells). This bias was driven by the anatomical distribution of TRN^SOM^-recipient neurons, which had a higher prevalence in the “core” (23.5%; 16 of 52 cells) compared to the “shell” (0%; 0 of 21 cells) in Som-Cre mice. In contrast, TRN^PV^-recipient neurons were equally likely to be found across the “core” (10.1%; 7 of 69 cells) and “shell” (10%; 1 of 10 cells) in PV-Cre mice. GPe→TRN inputs were equally likely to be found across the “core” (100%; 26 of 26 cells in Npas1-Cre mice) and the “shell” (100%; 5 of 5 cells in Npas1-Cre mice).

### Spatial and Kinetic Properties of Inhibitory Synaptic Inputs to TRN

Notably, extrinsic GPe→TRN inputs versus intra-TRN inputs differed in terms of their amplitude and kinetics. When comparing the kinetics of eIPSCs at the maximally active photostimulation site, eIPSCs evoked by GPe^Npas^ inputs were twofold larger in amplitude and two to threefold longer in half-width than those from TRN^PV^ or TRN^SOM^ neurons (**Figure 2K–L**). To investigate if these differences could be driven by the spatial location of GPe→TRN versus intra-TRN inputs (e.g., proximal or distal to the postsynaptic cell body), we turned to high-resolution focal photostimulation to map the afferent inputs.

**Figure 2.**
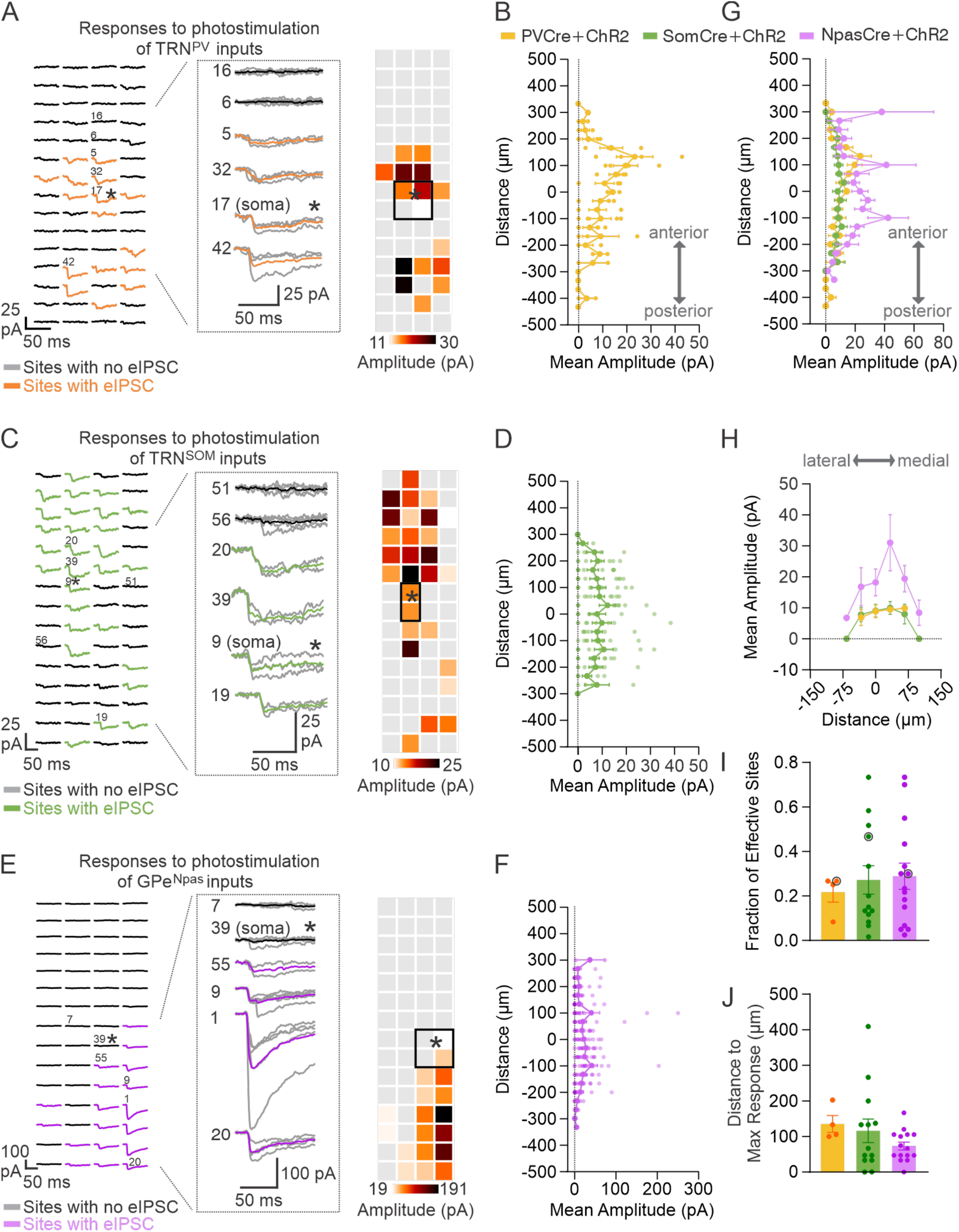
Spatial Distribution of Inhibitory Synaptic Inputs to TRN. **A.** Example photostimulation responses from a TRN^PV^-recipient TRN neuron obtained from a PV-Cre mouse expressing AAV2/5-DIO-ChR2-eYFP in TRN. Left: voltage-clamp recordings from a TRN cell held at **–**70 mV demonstrating the spatial distribution of monosynaptic GABAergic inputs upon optogenetic activation of PV+ inputs. The response map shows eIPSCs averaged across trials at each site between –20 and +200 ms relative to laser onset. A site was considered a “hot spot” (orange) if activation evoked eIPSCs ≥ 10 pA on ≥2 trials with event onset jitter ≤10 ms, with no spontaneous IPSC immediately preceding the laser onset. Nonresponsive sites (black traces) are averaged across all trials; “hot spots” (orange traces) are averaged across trials containing eIPSCs. Inset: sites labeled on the grid (left) are expanded (right), showing individual trials (grey) and the average (orange). Numbers indicate stimulation order. Right: heatmap of eIPSC amplitudes from 16 “hot spots” (of 60 sites), averaged across 6 trials. Grey indicates nonresponsive sites. The recorded cell body is indicated by the black square. **B.** Spatial distribution and mean amplitude of eIPSCs evoked by activation of *PV+* inputs along the anterior-posterior axis of the TRN. Data represent mean ± SEM from n = 4 responsive cells. **C–D.** Same as in (A**–**B), but from a TRN^SOM^-recipient TRN neuron recorded from a Som-Cre mouse expressing ChR2 in the TRN. “Hot spots” shown in green. Example heatmap shows eIPSCs from 28 “hot spots” averaged across 5 trials from one cell. Data in (D) represent mean ± SEM from n = 9 responsive cells. **E–F.** Same as in (A**–**B), but from a GPe^Npas^-recipient TRN neuron recorded from an Npas1-Cre mouse expressing ChR2 in GPe. “Hot spots” shown in purple. Example heatmap shows eIPSCs from 18 “hot spots” averaged across 5 trials from one cell. Data in (F) represent mean ± SEM from n = 14 responsive cells. **G.** Anterior-posterior distribution of the amplitude of eIPSCs mapped from PV, Som, or Npas inputs onto TRN neurons (mean ± SEM), averaged across rows using one map per cell. Distance is shown relative to the recorded cell. Maps were generated using a 15 × 4 grid (33 µm spacing) for 25 of 27 cells; two PV maps used 20 × 3 grids (33 µm spacing). PV maps: n = 4 cells; Som maps: n = 9 cells; Npas maps: n = 14 cells. **H.** Medial-lateral distribution of the amplitude of eIPSCs mapped from PV, Som, or Npas inputs onto TRN neurons, averaged across columns using one map per cell (mean ± SEM). **I.** Fraction of “hot spots” (photostimulation sites evoking synaptic responses), obtained by stimulating PV, Som, or Npas fibers in TRN. PV maps: n = 4 cells (3 mice); Som maps: n = 13 cells (9 mice), Npas maps: n = 15 cells (5 mice). One map from one cell was included in this analysis. Plots show mean ± SEM. Kruskal-Wallis test, not significant (*p* = 0.949; *p* = 0.96). **J.** Distance from recorded cell to site of maximal (trial-averaged) eIPSC amplitude in each map.

Both GPe^Npas^→TRN and intra-TRN–evoked IPSCs could be driven by activating sites up to 300 μm away from the postsynaptic cell body along the anterior–posterior axis, and up to ∼80 μm away along the medial–lateral axis (**Figure 2A–H**). Moreover, intra-TRN-recipient and GPe-recipient neurons exhibited a similar fraction of “effective” sites in the photostimulation protocol, indicating that both groups displayed a similarly high density of afferent connectivity (**Figure 2I**, **Supp. Figure 1C–D**).

For a subset of “responder” cells, we also mapped synaptic inputs with higher resolution closer to the cell body (66 × 66 μm grid) around the cell body (22 μm spacing), compared to a 460 × 100 μm grid (33 μm spacing). For brevity, we refer to these as microscale (<100 μm) and mesoscale (>100 μm) maps. We observed a striking difference across the distinct sources of inhibition when comparing synaptic responses evoked by microscale versus mesoscale inputs. Among GPe-recipient TRN responder cells, 90.9% (10 of 11 cells) displayed eIPSCs triggered by focal photostimulation using both micro- and meso-scale maps, suggesting that effective Npas1^+^ inputs were located both proximal and distal to the cell body. In contrast, only 28.5% of TRN^PV^-recipient neurons (2 of 7 cells) and TRN^SOM^-recipient neurons (4 of 14 cells) displayed eIPSCs triggered by focal photostimulation at both micro- and meso-scale maps. In other words, the majority of intra-TRN inhibitory connectivity was observed exclusively with meso-scale maps which primarily activate inputs distal to the cell body (71.5%; 6 of 21 cells), while GPe→TRN inhibitory connectivity was similarly recruited across micro- and meso-scale maps. Altogether, these results suggest that although intra-TRN recipient cells may be sparse, they display robust, wide-ranging afferent connectivity that is comparable to that observed in GPe^Npas^→TRN responder cells. However, GPe^Npas^ inputs provided stronger and longer-lasting inhibition in contrast to local TRN^PV^ and TRN^SOM^ inputs, demonstrating an intriguing dissociation between extrinsic versus local sources of inhibition.

### Rules of Intra-TRN Connectivity

TRN neurons are characterized by distinct molecular and electrophysiological hallmarks and are embedded in distinct neural circuits^8,9^. In light of this heterogeneity, we next investigated if subnetworks are preferentially engaged in recurrent inhibition within the TRN by leveraging the fact that PV and SOM are neurochemical markers of two major classes of TRN neurons^8,9,27^. Although we were unable to recover the neurochemical marker of each responder cell using immunohistochemistry, we used the presence of a direct ChR2-evoked photocurrent in the postsynaptic cell as a proxy for the whether the cell was transgene-positive or negative (i.e., TRN^PV+^, TRN^PV—^, TRN^SOM+^, or TRN^SOM—^ neurons). Based on prior literature, we assumed that Som-negative TRN cells are likely PV-positive^8,10^. ChR2-evoked photocurrents were distinguishable from synaptic responses due to their fast latency relative to laser onset and their persistence in the presence of gabazine in a subset of cells (**Supp. Figure 1B**).

In Som-Cre mice, only 1 of 17 TRN^SOM^-recipient TRN neurons exhibited direct photocurrents in response to photostimulation, suggesting that TRN^SOM+^ typically do not receive intra-TRN inputs from other TRN^SOM+^ neurons (i.e., reciprocal connections between TRN^SOM^ neurons are rare). Among the remaining 16 TRN^SOM^-recipient TRN neurons which failed to exhibit direct photocurrents, 100% were located in the TRN core (15 of 15 cells for which we were able to reconstruct the anatomical location). Given that the core and shell are predominantly populated by TRN^PV^ and TRN^SOM^ neurons, respectively,^8,10^ together with the absence of direct photocurrents in nearly all responders, we infer that the majority of TRN^SOM^-recipient TRN neurons have a high likelihood of being TRN^SOM—^. Taken together, these results reveal a dominance of TRN^SOM+^→TRN^SOM—^ connectivity (94.1% of responders) relative to TRN^SOM+^→TRN^SOM+^ connectivity (5.9% of responders). In PV-Cre mice, we classified 3 of 8 TRN^PV^-recipient TRN neurons as putative TRN^PV+^ neurons based on the presence of a direct photocurrent and anatomical location in the TRN core. These observations suggest a similar probability of TRN^PV+^→TRN^PV+^ connectivity (37.5% of responders) relative to TRN^PV+^→TRN^PV—^ connectivity (62.5% of responders), although generally, it was very rare for TRN^PV^ to be a presynaptic source of intra-TRN inhibition given the lower prevalence of responder cells when stimulating TRN^PV^ inputs (8.7%) compared to stimulating TRN^SOM^ inputs (18.6%). Taken together, these observations start to elucidate the rules of intra-TRN connectivity by establishing which pre-postsynaptic connections are possible.

### Feed-Forward Recruitment of Inhibition

The TRN is a critical component of corticothalamic feed-forward inhibition (CT→TRN→TC) and feed-back inhibition (TC→TRN→TC) of TC neurons. However, whether excitatory inputs to TRN can recruit intra-TRN inhibition in a feed-forward manner remains unknown. To this end, we injected ChR2-expressing AAV construct into the ventrobasal (VB) thalamus in adult mice under the Camk2α promoter, to specifically express ChR2 in excitatory glutamatergic TC neurons in VB thalamus as described^28^. Four weeks later, we prepared thalamic brain slices in which we severed the connection between TRN and VB to avoid circuit reverberation (feedback inhibition in the TRN→TC pathway that would cause rebound bursting in TC neurons, resulting in excitation of TRN)^28^. We then investigated the effect of optogenetic activation of ChR2-expressing TC axon terminals in TRN neurons (**Figure 3A–D**). Current-clamp recordings of tonic firing in TRN revealed a brief increase in firing upon TC terminal activation, as expected because TC neurons provide powerful excitatory input to TRN^28^. Excitation was followed by a pronounced interruption of firing lasting ∼200 ms before firing resumed (**Figure 3B, Supp. Figure 3**). This interruption was observed reliably across multiple trials in 21.7% of TRN neurons (**Figure 3C–D**) and was abolished by picrotoxin, confirming that it was mediated by GABA_A_ receptors. Furthermore, blocking AMPAR/NMDAR currents with kynurenic acid eliminated the initial firing rate increase caused by TC activation (**Figure 3B**). Because TC neurons do not project to GPe, these inhibitory effects must originate from intra-TRN connections. In summary, VB TC excitation can recruit GABAergic intra-TRN inhibition, establishing a feed-forward inhibitory motif within TRN.

**Figure 3.**
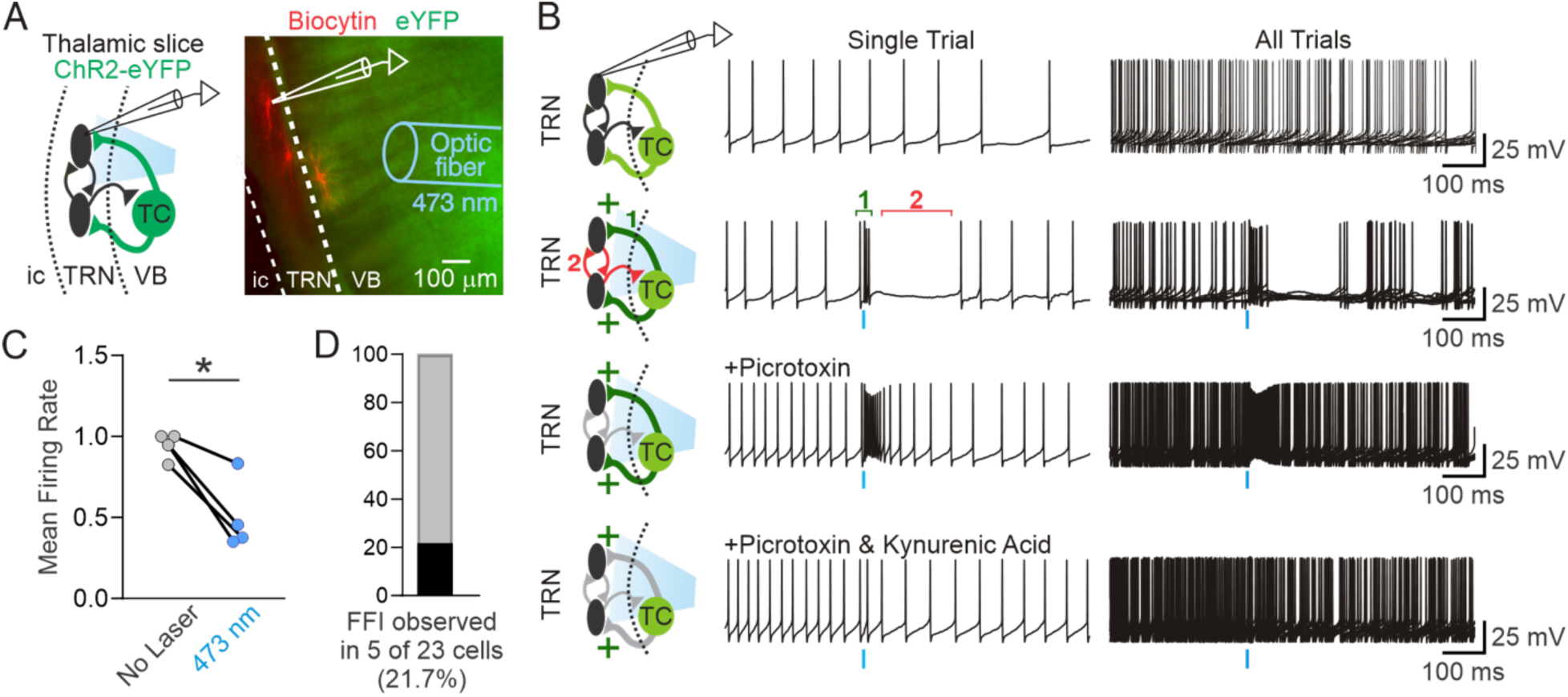
Feed-forward inhibition within the TRN generated by activation of ventrobasal thalamocortical axons in TRN. **A.** Schematic (left) and fluorescent image (right) showing ChR2 expression in thalamocortical (TC) neurons of the ventrobasal (VB) thalamus in adult mice. Whole-cell current-clamp recordings were performed from TRN neurons in horizontal thalamic slices, and 473 nm light pulses were delivered through an optical fiber positioned above the slice. Representative fluorescent image shows ChR2 (eYFP, green) expression and biocytin-filled TRN neurons (red). **B.** Example membrane potential trace from a TRN neuron. A 1 ms pulse of blue light evokes a transient depolarization followed by a pause in firing. **C.** Decrease in TRN firing rate relative to baseline following optogenetic activation of ChR2-expressing VB axons in TRN, indicating feed-forward inhibition. n = 4 cells. Data passed the Shapiro-Wilk normality test (*p* > 0.05); paired t-test on matched samples (*\*p* = 0.02). **D.** Proportion of TRN neurons in which TC-TRN-TRN feed-forward inhibition was observed (n = 5 of 23 cells, 4 mice).

## Discussion

### Dual inhibitory architecture of TRN: shedding light on the “searchlight” hypothesis

Together, these experiments reveal a dual inhibitory architecture of the TRN: a robust extrinsic input from GPe^Npas1^ neurons providing strong, uniform inhibition across the nucleus, and sparse recurrent inhibition from TRN^PV^ and TRN^SOM^ neurons which displayed spatial selectivity and could be recruited in a feed-forward manner by thalamocortical excitation. Our findings also revealed a potential dissociation between the postsynaptic locations of GPe versus intra-TRN inhibitory inputs. Using focal photostimulation to map connectivity at the micro- and meso-scale in the same neurons, we found that GPe-recipient TRN neurons exhibited synaptic responses at both scales, consistent with the presence of inhibitory inputs at both near-somatic and dendritic locations. In contrast, the relative sparsity of intra-TRN inputs evoked by micro-scale mapping suggests that their recruitment is biased towards more distal dendritic regions. While we did not perform ultrastructural analyses or reconstruct dendritic morphology, we speculate that intra-TRN inputs preferentially target distal dendrites while GPe inputs are recruited uniformly across the perisomatic and dendritic domains. These patterns suggest complementary modes of synaptic control by distinct inhibitory microcircuit motifs: extrinsic GPe^Npas^ inputs may be well-positioned to exert powerful, immediate inhibition at the soma, while recurrent intra-TRN inputs may shape integration through distal dendritic inhibition.

Crick’s “searchlight hypothesis” proposed that the TRN controls an internal attentional spotlight, amplifying select thalamocortical circuits through heightened burst firing in a subset of TRN neurons (Crick, 1984). Our findings refine this model by proposing two complementary modes of controlling the searchlight: pallidal inputs may serve as a strong, global coordinating signal that can rapidly shift TRN excitability (adjusting the “brightness” of the searchlight) while intra-TRN connections may serve as mediators of structured, selective interactions (adjusting the focus and location of the searchlight). Such parallel organization resembles motifs elsewhere in the brain, where local and long-range inhibition interact to balance stability and flexibility. For example, in hippocampal circuits, PV-expressing basket cells provide strong peri-somatic inhibition while long-range projections from oriens-lacunosum moleculare (O-LM) cells inhibit distal dendrites to gate excitatory inputs^29,30^. Future work should examine how intra-TRN and GPe inputs are coordinated during behavior, and whether they dominate under different brain states such as sleep, attention, or motor control^5^.

### Resolving intra-TRN inhibition: sparse nodes of convergence

Our results provide direct functional evidence and detailed characterization of intra-TRN chemical GABAergic synapses in adult mice, helping to resolve decades of conflicting reports obtained in a variety of species. Early studies reported reciprocal GABAergic connections between TRN neurons (e.g., see ^16,18^ and more recently^19^, while others, mainly in mice, failed to detect them^20–22^. In our hands, photostimulation of ChR2-expressing TRN^PV^ neurons recruited inhibition in only 8.6% of TRN neurons when using focal grid stimulation (estimated 1 μm spot size), but nearly 50% when using wider-field photostimulation (200 μm), suggesting that intra-TRN synaptic connections are distal and easily severed in slice preparations. These features would explain why paired recordings—which requires both pre- and post-synaptic cell bodies to be located in the same slice—or glutamate uncaging often failed to detect intra-TRN chemical inhibition, yet gap-junction mediated electrical synapses are routinely detected^14,15,20^ presumably because they are located more proximally along dendrites than chemical synapses. Further corroborating the sparse nature of intra-TRN synaptic connections, a recent study reported a single pair of TRN neurons coupled via GABAergic synapses amidst hundreds of paired recordings among closely apposed somata in the rat TRN^31^. Optogenetics circumvents this limitation by activating terminals even when the originating soma are absent, although opsin expression in axons remains a caveat in that low expression could lead to under-reporting the true prevalence. Taken together with previous reports that chemical synapses in the TRN are enriched along the anterior-posterior axis and follow distinct rules from electrical synapses^16^, our findings support the notion that intra-TRN connectivity is structured rather than uniform, and likely tailored to specific computational roles.

Our experiments revealed that although they may represent a small fraction of the TRN, intra-TRN recipient cells nevertheless exhibit wide-ranging afferent connectivity. The sparse but broadly connected nature of intra-TRN motifs suggests an intriguing, inverse analogy to “hub cells” described in the hippocampus, where a sparse set of inhibitory interneurons exert disproportional influence by broadcasting inhibition across a broad network ^32^. Intra-TRN recipient cells may represent nodes of convergence: although sparse, these TRN “convergence” cells are well-positioned to gather inhibitory inputs which may be spatially distributed across distinct TRN sectors and/or modalities, and subsequently gate TRN output in a state-dependent manner.

### Implications for TRN function

Our demonstration of the relative abundance of intra-TRN GABAergic synapses, compared to previous studies, expands the computational repertoire of the TRN. By providing localized, dendro-dendritic inhibition, intra-TRN synapses may sharpen receptive fields, modulate synchrony, and enable flexible attentional gating^4,33^. Their organization complements electrical synapses, which synchronize activity within ∼40 μm and occur in ∼30% of TRN neurons^15,34^. Whereas electrical coupling synchronizes neurons within the same domain, chemical inhibition may mediate lateral competition between adjacent domains or across modalities^17^. For instance, lateral inhibition stabilizes one and two-dimensional attractor dynamics, giving rise to head-direction selectivity in the anterior thalamus (reviewed by ^35^) and the characteristic hexagonal firing patterns of grid cells in the medial entorhinal cortex as rodents navigate their environments^36^. The recurrent inhibition within the TRN described in this study thus represents a novel circuit mechanism which may endow the TRN with hitherto undescribed computational motifs.

Our study focused on two major subnetworks in the TRN using PV and SOM as markers. While there are several other ways to parcellate TRN subpopulations using other neurochemical markers (e.g., the expression of Calbindin versus Somatostatin, or Spp1 versus Ecel1^10,11^, they generally map onto similar delineations of TRN subnetwork in terms of their electrophysiological properties and input-output connectivity. We thus consider our results to be generalizable, regardless of the nomenclature of choice. Importantly, we reveal a novel interaction between two subtypes of TRN neurons: although both PV and SOM neurons were capable of being pre- or post-synaptic partners of intra-TRN connections, (1) PV neurons were far less likely to serve as presynaptic sources of inhibition compared to SOM neurons, and (2) SOM neurons were more likely to inhibit PV neurons compared to other SOM neurons, while PV neurons inhibited other PV neurons and SOM neurons similarly. By revealing a previously undescribed recurrent inhibitory motif between TRN neurons embedded in distinct circuits, our findings position inhibitory microcircuits of the TRN as important contenders in shaping thalamocortical computations.

### Disease relevance

GABAergic inhibition between TRN cells has been proposed to play a key role in constraining thalamocortical synchrony^37^. Disruption of intra-TRN GABAergic transmission, whether by genetic deletion of GABA_A_ receptor β3 subunits or hemizygous loss of sodium channel encoding gene *Scn8a*, produces pathological oscillations and absence seizures^19,33,37^. Moreover, loss of GABAergic inhibition in TRN has been suggested to drive epileptiform discharges during sleep spindles after mild brain injury^38^. Given that intra-TRN chemical and electrical synapses are likely engaged differently across states, selective impairments could yield distinct circuit phenotypes. Our work also suggests that GPe may be a stronger modulator of TRN than previously appreciated, raising the possibility that dysfunction in the GPe^Npas^→TRN pathway could destabilize thalamocortical rhythms, contributing to movement disorders or attentional deficits. Beyond epilepsy, TRN dysfunction has been implicated in schizophrenia, Alzheimer’s disease, ADHD, and autism spectrum disorders, where sensory gating and attentional control are impaired^39,40^. Our work dissecting the cell-type specific inhibitory microcircuits in the TRN thus opens the door towards investigating specific inhibitory synapses in the TRN as potential cellular targets for therapeutic interventions.

### Limitations and future directions

Although the current study focused on GPe as an external source of GABAergic inhibition to TRN, there are other sources of inhibition (e.g., lateral hypothalamus^41^) for which the cell-type connectivity rules are unknown. Our experiments were confined to a single spatial plane, leaving unexplored potential differences along the anterior–posterior axis, which maps onto distinct sensory modalities. Such mapping could clarify whether intra-TRN inhibition mediates cross-modal gating, providing the synaptic substrate for Crick’s “guardian of the gateway” hypothesis^42^. Furthermore, our experiments revealed both high and low prevalence of intra-TRN inhibition depending the technique (wider-field illumination versus focal photostimulation). The true prevalence likely lies between these extremes and varies with behavioral state, and characterizing the extent intra-TRN recruitment in vivo will be a critical step towards illuminating the physiological relevance of recurrent inhibition within the TRN.

## Methods

We performed all experiments in accordance with protocols approved by the Institutional Animal Care and Use Committee at the University of California, San Francisco, and the Gladstone Institutes. All precautions were taken to minimize animal stress and reduce the number of animals used.

### Mice

We performed all experiments per protocols approved by the Institutional Animal Care and Use Committee at the University of California, San Francisco and Gladstone Institutes. Precautions were taken to minimize stress and the number of animals used in all experiments. Adult (P30– P180) male and female mice were used: wild-type C57BL/6J mice (wild-type, IMSR_JAX:000664; n = 6 mice), Somatostatin (SOM)-Cre mice (SOM-IRES-Cre, IMSR_018973; n = 17 mice), Parvalbumin (PV)-Cre mice (PV-Cre, IMSR_JAX: 017320; C57BL/6 congenic; n = 26 mice), Npas1-Cre-2A-tdTomato BAC transgenic line (NpasCre; RRID:ISMR_JAX:027718; gift from Dr. Aryn Gittis; n = 8 mice). Adult (P30–P180) male and female mice were used: wild-type C57BL/6J (IMSR_JAX:000664; n = 6), Somatostatin (SOM)-Cre (SOM-IRES-Cre, IMSR_018973; n = 17), Parvalbumin (PV)-Cre (IMSR_JAX:017320; C57BL/6 congenic; n = 26), and Npas1-Cre-2A-tdTomato BAC transgenic mice (Npas1-Cre; RRID:IMSR_JAX:027718; gift from Dr. Aryn Gittis; n = 8).

### Viral injections

We performed viral injections into TRN, VB thalamus, or GPe as described ^8,9,28,43,44^ in mice aged 2–7 months. Briefly, injections using a 10-μl syringe and 34-gauge needle were controlled by pump with a flow rate set to 100 nL/min (World Precision Instruments). **TRN:** 100 nL unilateral injections of AAV2/5-EF1a-DIO-hChR2(H134R)-EYFP (Addgene plasmid #20298; RRID:Addgene_20298) at stereotaxic coordinates 1.3–1.6 mm posterior to Bregma, 2.0–2.1 mm lateral, and 2.6–3.4 mm ventral to the cortical surface. **GPe:** 80 nL unilateral injections of the same virus in Npas-Cre mice at –0.28 mm posterior, 2.1 mm lateral, and 4.0 mm ventral from the cortical surface. **VB thalamus:** 200 nL unilateral injections at 1.3–1.6 mm posterior, 1.6–1.7 mm lateral, and 3.0–3.2 mm ventral to the cortical surface.

### Optogenetic stimulations for whole slice illumination experiments

Optogenetic responses were elicited by activating ChR2-expressing PV-Cre neurons with 450 nm light (OEM Laser Systems, 0.8–2 mW) delivered via a 200-μm optical fiber (Thorlabs) positioned over the TRN (**Supp. Figure 2**).

### Slice preparation

Thalamic slices were obtained as previously described^8,43^. Mice were euthanized with isoflurane and transcardially perfused with ice-cold sucrose solution (234 mM sucrose, 2.5 mM KCl, 1.25 mM NaH2PO4, 10 mM MgSO4, 0.5 mM CaCl2, 26 mM NaHCO3, and 11 mM glucose equilibrated with 95% O2 and 5% CO2). Horizontal 250-μm slices containing TRN were cut using a Leica VT1200 microtome (Leica Microsystems). Slices were incubated at 32 °C for 30 min, then recovered for at least 1 h at 24–26 °C in ACSF (126 mM NaCl, 2.5 mM KCl, 1.25 mM NaH₂PO₄, 2 mM MgCl₂, 2 mM CaCl₂, 26 mM NaHCO₃, 10 mM glucose) equilibrated with 95% O₂ / 5% CO₂.

### Patch-clamp electrophysiology

Recordings were performed as previously described^8,43^. TRN neurons were visually identified by differential interference contrast (DIC) optics (Olympus microscope, 63× objective, NA 1.1, WD 1.5 mm; SKU 1-U2M592). Signals were digitized using Axon Digidata 1550 and pClamp 10, and amplified with MultiClamp 700B (Molecular Devices). Recording electrodes (borosilicate glass, Sutter Instruments) had resistances of 2.5–4 MΩ when filled with intracellular solution. Cells were included for analysis if the access resistance was < 25 MΩ, with the exception of one GPe→TRN recipient cell (35 MΩ). Access resistance did not differ between responders (15.6 ± 4.9 MΩ, n = 62 cells) and non-responders (15.9 ± 3.9 MΩ, n = 157 cells; unpaired t-test, *p* = 0.68). For mapping experiments, the internal solution contained 135 mM CsCl, 10 mM HEPES, 10 mM EGTA, 2 mM MgCl₂—6H₂O, and 5 mM QX-314 (pH adjusted to 7.3 with CsOH, 290 mOsm). Recordings were performed at room temperature in the presence of kynurenic acid (2 mM; Sigma-Aldrich #K3375) in ACSF. To block GABA_A_R-mediated currents, gabazine (GBZ; 50 μM; Sigma #SR-95531) prepared in dimethyl sulfoxide (DMSO; Sigma #D8418) was bath-applied.

### LASU photostimulation

Optogenetic mapping was performed using the LASU (Laser Applied Stimulation and Uncaging) system (Scientifica, Ltd). This custom-made system includes an epifluorescence SliceScope microscope (Scientifica) with 4× and 63× water-immersion objectives (Olympus) and a uEye camera (IDS). Focal photostimulation used a 405 nm laser. For photostimulation at multiple sites within the field of view, the laser spot was repositioned using a set of two galvanometer scan mirrors controlled by LASU software (v24.1). The laser spot size is estimated to be approximately 1 μm diameter at 63×, and 10 μm diameter at 4×. Laser power, which is controlled by the LASU software, was determined using a power meter, and ranged 0.1–30 mW, with lower powers for GPe→TRN maps (mean 0.4 mW) and higher for intra-TRN maps (mean 3.65 mW). Photostimulation parameters (location, duration, frequency) were defined and saved in LASU software for offline analysis.

#### Post-hoc imaging

Fixed slices (4% PFA, then 30% sucrose) were imaged using a Zeiss Axio Examiner A1 microscope (GFP and brightfield) to confirm ChR2-EYFP expression. Anatomical landmarks were used to align fixed and live slice images containing the stimulation grid. High-resolution images from Zen software (Zeiss) were used to estimate actual grid dimensions generated by LASU software (LASU v2.1). Specifically, 5 slices with clear alignment between fixed and live slice images were used to empirically determine scaling factors, which ranged between 55.5% and 70.1%. To ensure a conservative estimation, the lowest scaling factor was used (55.5%) to rescale distance across all experiments. For example, a grid nominally 840 × 180 μm (15 × 4 sites, 60 μm spacing) corresponded to a true span of 466.2 × 99.9 μm (33.3 μm spacing) after scaling.

#### Mapping synaptic connectivity

Voltage-clamp recordings were obtained in Clampex (Molecular Devices) and controlled via LASU software (v24.1). For each cell, mapping was first performed at high magnification (63x objective) to determine the presence or absence of photocurrents and the optimal laser power. Stimulation parameters and grid properties were generated in the LASU software (v24.1). 63x stimulation typically consisted of 160 x 160 μm grids (40 μm spacing) centered around the soma. After determining responses at 63x, the objective was switched to 4x to test connectivity from a larger region of the TRN. 4x stimulation typically consisted of 900 x 240 μm grids (60 μm spacing), although in some cases, the grid spacing and/or shape had to be modified depending on the curvature and thickness of the TRN in the slice. Laser stimulation consisted of 5 ms pulses at each spot, with an inter-pulse-interval of 500-ms (2-Hz) separating consecutive spots, unless otherwise specified. Each trial consisted of each spot being stimulated in a random order; the random order was preserved across trials for each grid. Re-randomization of the stimulation order within the same grid resulted in nearly identical response maps. Coordinates for each point on the stimulation grid, as well as images of the stimulation grid on the slice, were saved for offline analysis. The position of the recorded cell and stimulation grid were reconstructed offline using tiled images of the recorded slice which were cross-registered with higher-resolution bright field and fluorescent images obtained from a Zeiss Axio A1 epifluorescent scope.

#### Generation of response maps and kinetic analysis of synaptic responses

Response maps were built in Python using the pyABF package and SciPy library. Voltage-clamp recordings obtained in gap-free mode were binned into 520-ms sweeps (including 20-ms preceding the laser pulse) and low-pass Bessel filtered at 500 Hz, with each sweep corresponding to a photostimulation site in a given trial. Sweeps were assigned to the corresponding photostimulation site using the grid coordinates associated with each experiment, which were then used to generate the grid of voltage-clamp responses (e.g., **Figure 2**). The presence of eIPSCs was determined for each photostimulation site and trial (for each experiment) by two investigators. The inclusion criteria for eIPSCs was as follows: an inward (negative) current with amplitude greater than 10 pA, asymmetric rise and decay (i.e., fast rise, slow decay), occurring after 5 ms from laser onset and within 50 ms from laser offset. A given photostimulation site was considered a “hot spot” if it contained at least 2 trials with an evoked synaptic current with consistent onset latency (within 10 ms of each other) and did not contain a spontaneously occurring IPSC preceding the laser onset. This manual curation was used to generate a “masking file” for each experiment, which was used in conjunction with the SciPy find_peaks function, to identify the amplitude of synaptic currents at photostimulation sites and trials containing eIPSCs. The find_peaks function (parameters: prominence=2, threshmax=10000) was used to extract only the amplitude, which was visually cross-validated against raw traces. The orientation of response maps were aligned to the mouse brain atlas (anterior/posterior, medial/lateral). The distance between the stimulation site and somatic recording site was estimated by approximating the location of the recording pipette tip on the stimulation grid to within 2 to 4 sites. eIPSCs identified above were analyzed separately in Clampfit (Molecular Devices) to automatically extract half-width, decay time, area and amplitude, and manually extract rise time and latency. All events imported into Clampfit were manually validated to be eIPSCs to exclude spontaneously occurring IPSCs or artifacts. Kinetic analyses presented in **Figure 1** include eIPSCs observed at the maximally active photostimulation site for a given cell, while analyses presented in **Figure 2** include all eIPSCs detected across all photostimulation sites. In **Figure 2I** and **Supp. Figure 2C**, comparison was performed only for responder cells which were mapped at low magnification. Maps were typically obtained using a 15 x 4 grid (60 μm spacing).

#### Current-clamp analysis

Data were analyzed in Clampfit (Molecular Devices). We determined the following epochs per trial for analysis: baseline (200 ms preceding laser pulse), laser (30ms starting at laser onset for square pulses, and 100 ms for ramp pulses), post-laser (200 ms following laser epoch), and recovery (200 ms following post-laser). Current injection duration was 2 s. Mean firing rates were calculated per epoch (spikes / epoch duration) and averaged across trials. Firing-rate histograms (**Figure 2E**) were binned at 10 ms and normalized by trial count. For statistical analyses (**Figure 2G–H**), only matched samples were included. Each cell’s mean firing rate (or baseline-normalized rate) in laser vs. no-laser conditions was compared using paired or ratio-paired t-tests.

## Acknowledgements

We would like to thank Dr. Aryn Gittis and Dr. Savio Chan for Npas1-Cre mice, Irene Lew for help with animal husbandry, and Dr. Savio Chan and Isaac Chang for insightful discussions about Npas1-Cre mice.

## Funding

FSC was supported by NINDS F31 NS111819; JRH was funded by NINDS NS34774; JTP was supported by the Gladstone Institute of Neurological Disease.

## Author Contributions

Conceptualization: JTP, JRH. Methodology: FSC, JRH, JTP. Investigation: FSC, JTP. Data curation: FSC, JTP. Analysis: FSC, JTP. Visualization: FSC, JTP, Supervision: JTP, JRH. Funding acquisition: FSC, JRH, JTP. Writing – original draft: FSC, JTP. Writing – reviewing & editing: all authors.

---

**Supplemental Figure 1.**
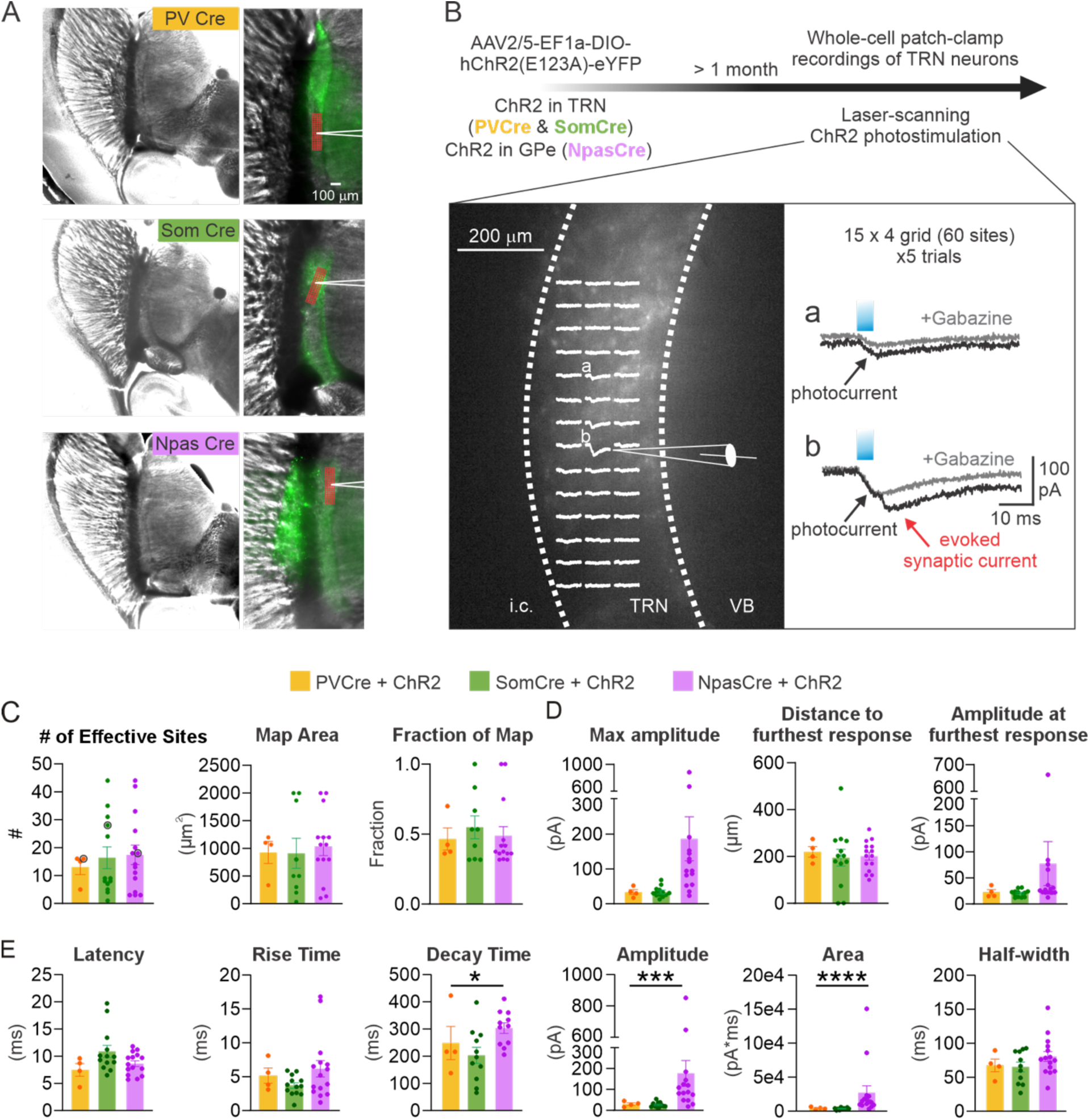
Characterization of optogenetic mapping and spatial organization of GABAergic inputs to TRN neurons. Related to Figures 1, 2. **A.** Brightfield and fluorescent images of horizontal thalamic slices (2.5x magnification, left; 5x magnification, right) from PV-Cre and Som-Cre mice injected with AAV2/5-DIO-ChR2-eYFP (green) into TRN, and from an Npas-Cre mouse injected with the same viral construct into GPe. Right: example showing the position of the recording pipette and the optogenetic stimulation grid (red). **B.** Overview of the photostimulation experiment. Inset shows a brightfield image of TRN expressing ChR2 in a PV-Cre mouse. Average responses from focal photostimulation at each site in the 15 x 4 grid are overlaid on the slice image from one recorded TRN neuron. (a) indicates a site evoking only a photocurrent, visible immediately upon the laser pulse and unchanged by Gabazine application. (b) indicates a site evoking both a photocurrent and a delayed inward current (eIPSC), which is selectively abolished by Gabazine. **C.** Extent of synaptic connectivity from neurochemically and anatomically distinct sources of GABAergic input. Number of “hot spots” (effective photostimulation sites) obtained by stimulating TRN^PV^, TRN^SOM^, or GPe^Npas^ fibers onto postsynaptic TRN neurons (left). PV maps: n = 4 cells (3 mice); Som maps: n = 13 cells (9 mice), Npas maps: n = 15 cells (5 mice). One map from one cell was included in this analysis. Bar graphs show mean ± SEM. Kruskal–Wallis test, not significant (*p* = 0.949; *p* = 0.96). Area representing the anterior–posterior and medial–lateral extent of responses (“map area”) (center) and the fraction of responsive sites within this area (right). **D.** Amplitude of the site of maximal, trial-averaged eIPSC response (left), distance from the recorded cell to the most distal eIPSC response (center), and trial-averaged eIPSC amplitude at that site (right). **E.** Properties of eIPSCs recorded at the maximal responsive site for each map. PV: n = 4 cells; Som: n = 13 cells; Npas: n = 15 cells. **Kruskal–Wallis test, *p* < 0.05, \*\**p* < 0.0005, \**p* < 0.00005.

**Supplemental Figure 2.**
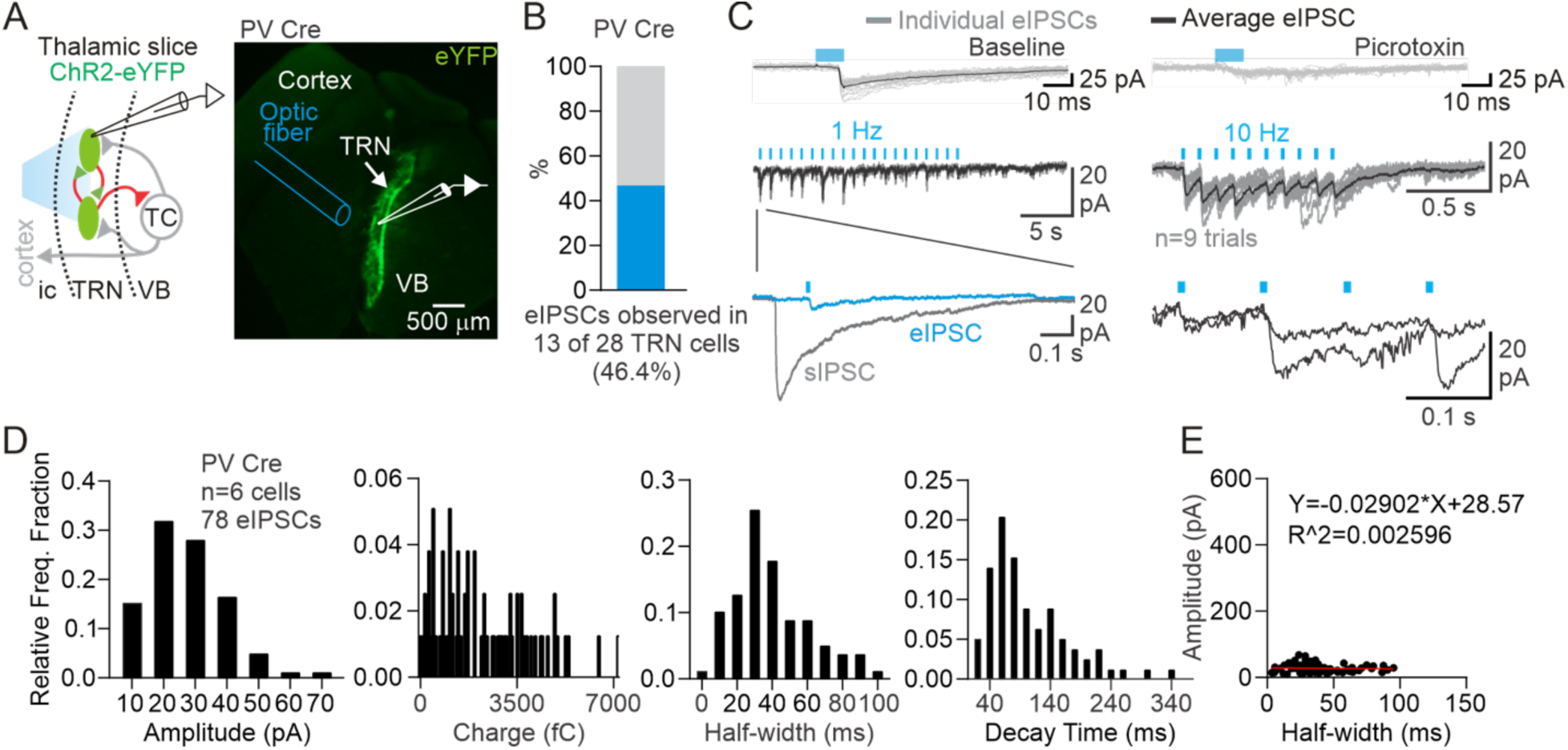
Optogenetic activation of intra-TRN GABAergic synapses in PV-Cre mice. Related to Figure 1. **A.** Schematic (left) and fluorescent image (right) of a horizontal thalamic slice from a PV-Cre mouse expressing ChR2 in TRN (AAV injection into the TRN). Whole-cell voltage-clamp recordings were performed from TRN neurons, while an optical fiber delivered 473 nm light pulses to activate ChR2- or ChETA-expressing axons (not shown). **B.** Top: example synaptic currents evoked by ChR2 activation, blocked by picrotoxin. Middle: eIPSCs reliably entrained at 1 Hz and 10 Hz stimulation. **C.** Fraction of TRN cells showing eIPSCs following wider-field (200 um) photoactivation of TRN^PV^ inputs (n = 13 of 28 cells; ChETA). **D.** Kinetic properties of eIPSCs evoked by TRN^PV^ activation. Distributions of amplitude, charge, half-width, and decay time (n = 78 eIPSCs from 6 cells). **E.** Relationship between eIPSC half-width and amplitude.

**Supplemental Figure 3.**
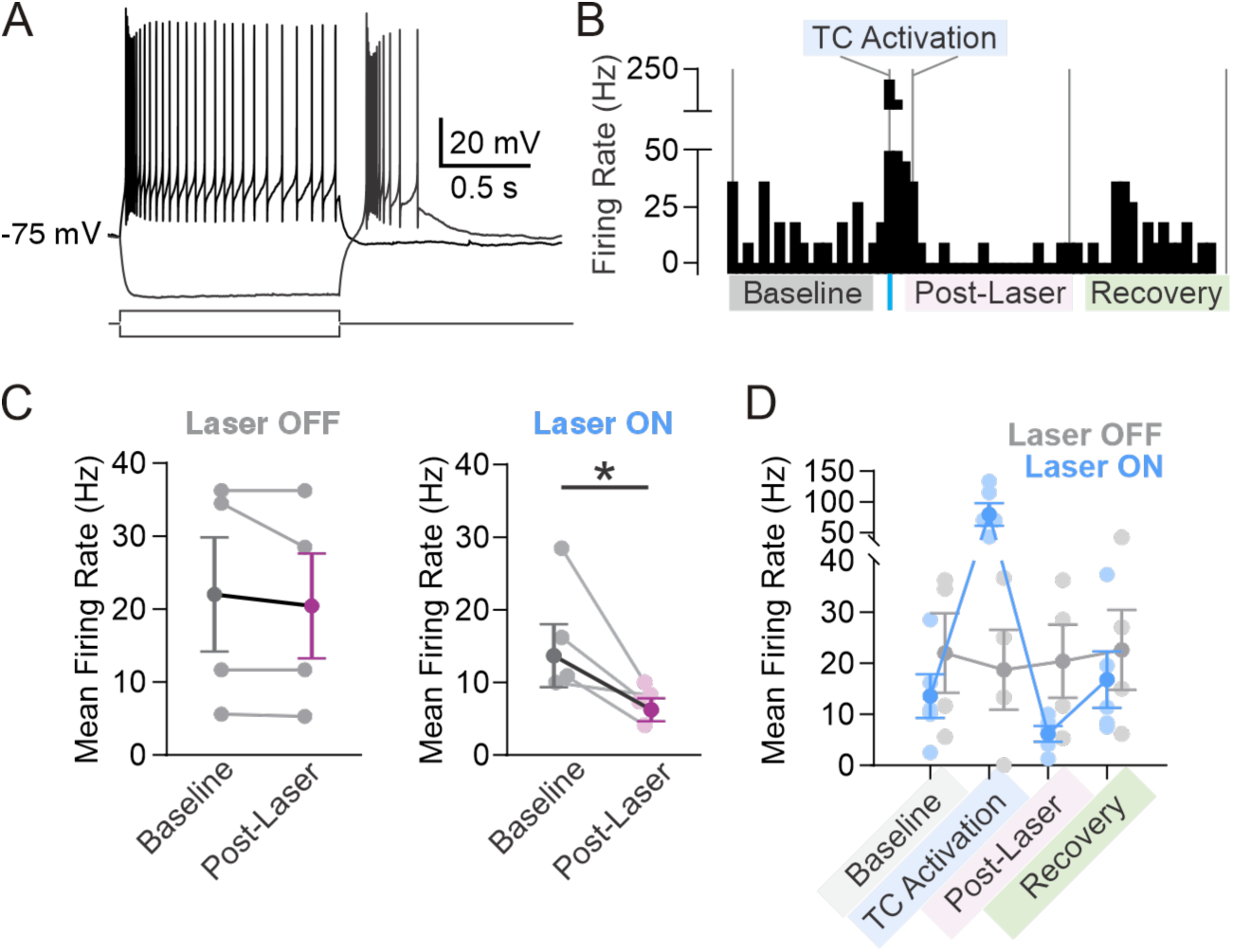
Optogenetic activation transiently suppresses TRN neuron firing. Related to Figure 3. **A.** Representative whole-cell current-clamp traces of a TRN neuron in response to depolarizing and hyperpolarizing current steps. **B.** Firing rate of example TRN neuron exhibiting feed-forward inhibition upon activation of ventrobasal thalamocortical axons (same as in **Figure 3B**). A 1 ms blue light pulse triggers a brief increase followed by suppression of firing. Vertical lines indicate analysis epochs: baseline (200 ms), laser (30 ms), post-laser (200 ms), and recovery (200 ms). **C.** Light-evoked suppression of firing rate observed with ChR2-activation (right) but not during no-laser control trials (left). n = 4 cells. Data passed the Shapiro–Wilk normality test (*p* > 0.05). Ratio paired t-test: no laser, *p* = 0.27 (ns); 473 nm, *\*p* = 0.032. Same data shown in Figure 3C. **D.** Mean firing rate across epochs in responder cells (n = 5) with (blue) and without (grey) laser stimulation. Two-way repeated-measures ANOVA: interaction, \*\*\**p* < 0.0001; time, \**p* = 0.0121; laser, *p* = 0.21 (ns); subject, *p* = 0.21 (ns).

## Notes

### Competing Interest Statement

The authors have declared no competing interest.

